# ADAR2 mediated Q/R editing of GluK2 regulates homeostatic plasticity of kainate receptors

**DOI:** 10.1101/308650

**Authors:** Sonam Gurung, Ashley J. Evans, Kevin A. Wilkinson, Jeremy M. Henley

## Abstract

Kainate receptors (KARs) are heteromeric glutamate-gated ion channels that regulate neuronal excitability and network function in the brain. Most KARs contain the subunit GluK2 and the precise properties of these GluK2-containing KARs are determined by additional factors including ADAR2-mediated mRNA editing of a single codon that changes a genomically encoded glutamine (Q) to arginine (R) in the pore-lining region of GluK2. ADAR2-dependent Q/R editing of GluK2 is dynamically regulated during homeostatic plasticity (scaling) elicited by suppression of synaptic activity with TTX. Here we show that TTX decreases levels of ADAR2 by enhancing its proteasomal degradation. This selectively reduces the numbers of GluK2 subunits that are edited and increases the surface expression of GluK2-containing KARs. Furthermore, we show that partial ADAR2 knockdown phenocopies and occludes TTX-induced scaling of KARs. These data indicate that activity-dependent regulation of ADAR2 proteostasis and GluK2 Q/R editing provides a mechanism for KAR homeostatic plasticity.

## Introduction

Kainate receptors (KARs) are ionotropic glutamate receptors that are tetrameric assemblies composed of combinations of five receptor subunits, GluK1-5 (1, 2). KARs can be located pre-, post– and/or extrasynaptically, where they contribute to neurotransmitter release, postsynaptic depolarisation and the regulation of neuronal excitability (3). The variety of possible subunit combinations and co-assembly with Neto auxiliary subunits (4), creates a wide range of possible KAR subtypes, with receptors containing GluK2 and GluK5 being the most abundant (3).

Additional diversity of GluK2 containing KARs arises from RNA editing of GluK2 (5, 6) by the nuclear ADAR2 enzyme which edits pre-mRNAs encoding GluK2 and GluK1, as well as the AMPAR subunit GluA2, and other non-coding RNAs (7, 8). ADAR2 mediated Q/R editing in the pore-lining region of GluK2 alters a genomically encoded glutamine residue to an arginine, changing both the calcium permeability and biophysical properties of the KARs. More specifically, GluK2(R) shows markedly reduced tetramerisation leading to its accumulation in the ER (9). Furthermore, assembled GluK2(R)-containing KARs that do reach the plasma membrane are calcium impermeable and have a channel conductance of less than 1% of non-edited GluK2(Q)-containing KARs (10).

ADAR2 levels are very low during embryogenesis but increase in the first postnatal week (11) to edit ~80% of GluK2, ~40% of GluK1 and ~99% of GluA2 subunits in the mature brain (12–14). ADAR2 knockout mice die at the early postnatal stage, but can be rescued by expressing the edited form of GluA2, demonstrating that unedited AMPARs are fatally excitotoxic (15). In contrast, mice specifically deficient in GluK2 Q/R editing are viable but are seizure prone and adults display an aberrant form of NMDAR-independent long term potentiation (LTP) (16), demonstrating the importance of GluK2 editing to network function and synaptic plasticity.

NMDARs, AMPARs and KARs have all been shown to play roles in synaptic plasticity and their dysregulation is a prominent feature of cognitive decline in aging and neurodegenerative diseases (17). LTP and LTD (long-term depression) enhance or decrease the efficiency of synaptic transmission, respectively, and are primarily mediated by changes in the number and properties of AMPARs at the postsynaptic membrane. ‘Classical’ LTP requires the activation of NMDARs (18, 19). However, recent data from our lab has demonstrated a novel role for KARs as critical inducers of one form of LTP (KAR-LTP_AMPAR_) (20). Moreover, in addition to directly inducing synaptic plasticity of AMPARs, KARs themselves are plastic and undergo LTD (21) and LTP (22–24).

The assembly and progression of GluK2-containing KARs through the secretory pathway is controlled by a series of activity regulated checkpoints (25). Furthermore, chronic suppression of synaptic activity with TTX ‘up-scales’ KAR surface expression (25). Homeostatic plasticity (scaling) of AMPARs and NMDARs is essential for normal neuronal network and brain function because it constrains neuronal firing to within a tunable physiological range (26–28). The discovery of KAR scaling is important because regulation of the number of surface expressed KARs will be a critical factor for induction of KAR-LTP_AMPAR_, and therefore constitutes a key regulator of synaptic transmission and neuronal excitability. However, how KAR scaling occurs mechanistically is unclear.

Here we show that chronic suppression of network activity with TTX leads to the proteosomal degradation of ADAR2. This, in turn, reduces editing of GluK2 pre-mRNA with a consequent increase in unedited GluK2, thus enhancing homomeric KAR assembly, ER release and surface expression. Importantly, this up-scaling mechanism is specific to GluK2 as TTX did not change the editing status of GluA2. Together these data demonstrate a novel and selective role of mRNA editing by ADAR2 in homeostatic plasticity of KARs.

## Results

### Chronic suppression of network activity with TTX increases surface levels of GluK2-containing KARs

Chronic TTX treatment increases AMPAR surface expression and AMPAR excitatory postsynaptic currents (EPSCs) (reviewed in (29)). Surface levels of GluK2 containing KARs also upscale in response to TTX treatment (25). Consistent with this, we show robust scaling of both KAR (Figure 1A,B) and AMPAR subunits (Figure 1A,C). In contrast, epidermal growth factor receptor (EGFR), which is widely expressed at synapses (30), was not affected by TTX (Figure 1A,D). Intriguingly, TTX-mediated suppression of synaptic activity treatment also reduced GluK2 Q/R editing (Figure 1E,F) but did not affect Q/R editing of the AMPAR subunit GluA2 (SI Figure 1A,B,C). These findings suggest that KAR, but not AMPAR, up-scaling by TTX treatment may be mediated by decreased levels of GluK2 Q/R editing.

**Figure 1:**
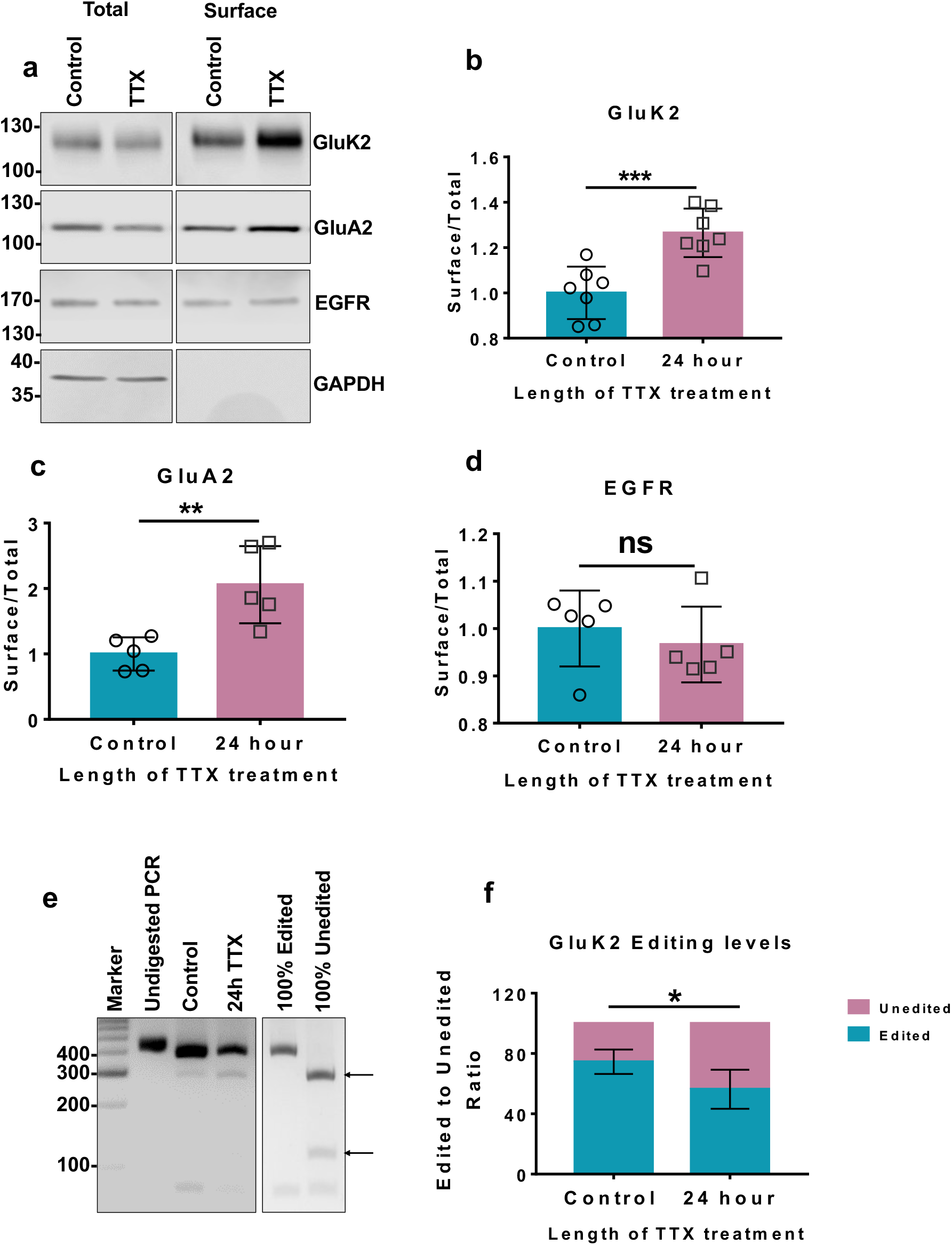
Chronic suppression of network activity with TTX increases surface levels of GluK2 containing KARs. a. Representative western blot of total and surface levels of GluK2 (KAR subunit), GluA2 (AMPAR subunit), EGFR and GAPDH in hippocampal neurones with or without 24 h TTX treatment to suppress synaptic activity. EGFR was used as negative control and GAPDH was used as a control to show only surface proteins were labelled with biotin. b. Quantification of surface levels of GluK2 (A) from 5 independent experiments. Surface levels were normalised to their total levels. Statistical Analysis; Unpaired t-test ***<0.001. c. Quantification of surface levels of GluA2 (A) from 5 independent experiments. Surface levels were normalised to their total levels. Statistical Analysis; Unpaired t-test **<0.01. d. Quantification of surface levels of EGFR (A) from 5 independent experiments. Surface levels were normalised to their total levels. Statistical Analysis; Unpaired t-test, ns>0.05. e. RT-PCR and BbvI digestion analysis of GluK2 Q/R editing from hippocampal neurones treated with or without TTX. f. Quantification of E from 5 independent experiments. Statistical Analysis: Unpaired t-test; *<0.05.

### Chronic suppression of network activity with TTX decreases ADAR2 levels

We next tested the effects of chronic suppression of network activity on ADAR2 levels. Incubation with TTX for 24 h decreased ADAR2 levels by ~50%, but ADAR1 levels were unaffected (Figure 2A-D). Interestingly, longer periods of TTX treatment did not decrease ADAR2 levels any further (Figure 2E,F), suggesting that a basal level of ADAR2 is retained even under long-term suppression of synaptic activity. As expected, the decrease in ADAR2 following 24 h TTX treatment occurs in the nucleus (Figure 2G,H), where ADAR2 binds to the GluK2 pre-mRNA prior to mRNA splicing and maturation (31). Moreover, both the intensity of ADAR2 signal and the percentage of ADAR2 expressing cells decreased significantly following 24 h TTX treatment (Figure 2I,J,K).

**Figure 2:**
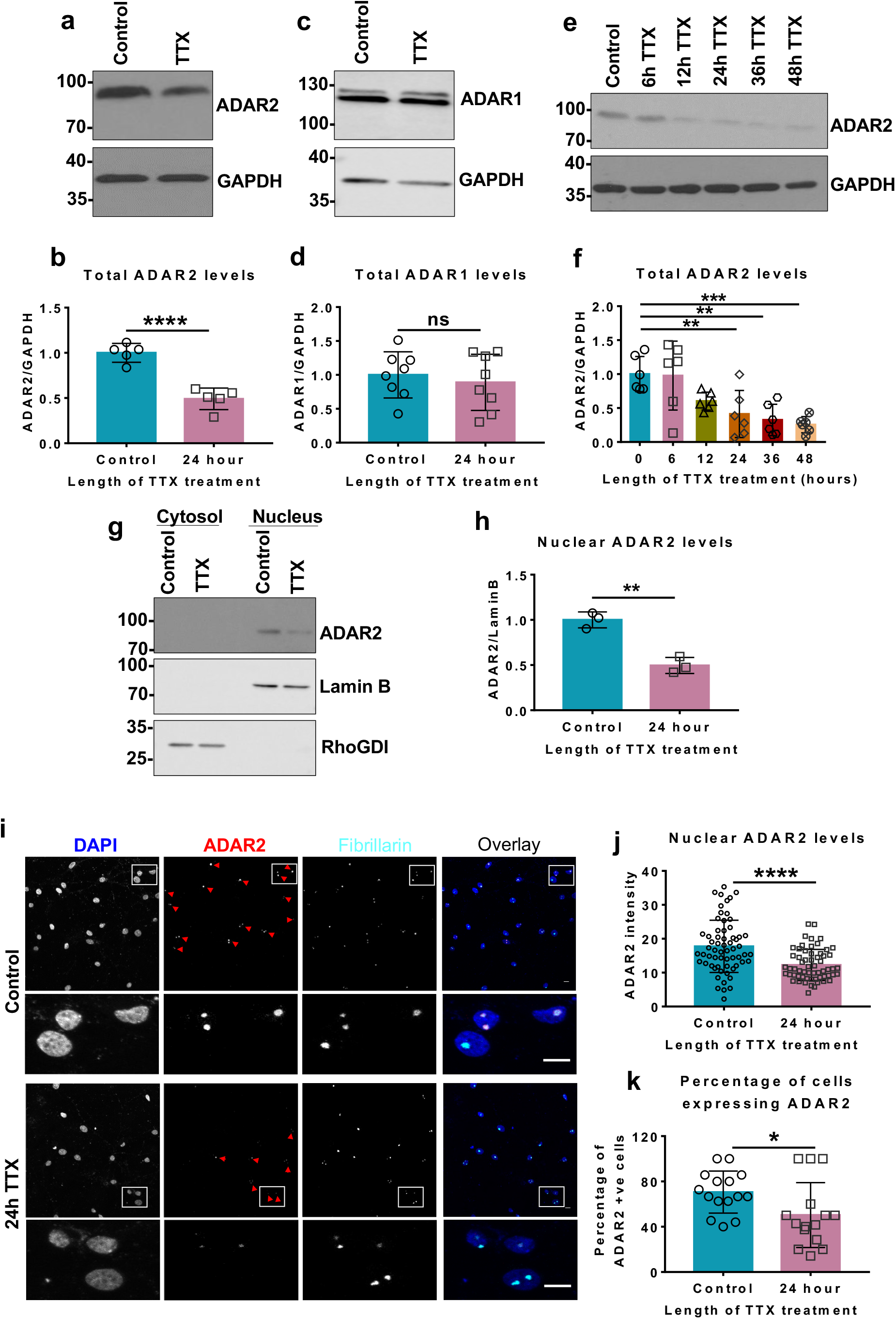
Chronic suppression of network activity with TTX decreases ADAR2 levels. a. Representative western blots of total ADAR2 and GAPDH levels in hippocampal neurons with or without 24 h TTX treatment to suppress synaptic activity. b. Quantification of (A) total ADAR2 normalised to GAPDH from 5 independent experiments. ADAR2 levels normalised to loading control GAPDH. Statistical Analysis: Unpaired T test; ****<0.0001. c. Representative western blots of total ADAR1 level and GAPDH levels in hippocampal neurones with or without 24 h TTX treatment. d. Quantification of (C) total ADAR1 normalised to GAPDH from 8 independent experiments. Both bands were quantified. Statistical Analysis: Unpaired T test; ns>0.05. e. Representative western blots showing total ADAR2 and GAPDH levels with increasing lengths of TTX treatment. f. Quantification of (E) total ADAR2 normalised to GAPDH from 6 independent experiments. Statistical Analysis: One Way ANOVA Dunnett’s multiple comparison test; **<0.01, ***<0.001. g. Representative western blot of nuclear ADAR2 levels in hippocampal neurones with or without 24 h TTX treatment. Cell fractionation was performed to determine the ADAR2 levels in the nucleus. Lamin B was used as a nuclear marker and RhoGDI as cytosol marker. h. Quantification of (G) nuclear ADAR2 immunoblots normalised to Lamin B from 3 independent experiments. Statistical Analysis: Unpaired T test; **<0.01. i. Representative images of hippocampal neurons with or without 24 h TTX treatment labelled with nuclear DAPI stain (blue), anti-ADAR2 (red) and anti-Fibrillarin (nucleolar marker; cyan). Bottom panels show zoom in images as indicated and the red arrows indicate cells expressing ADAR2. j. Quantification of (I) ADAR2 intensity per nucleus. N=3 independent dissections and n=60 cells for control and 62 cells for TTX treated. Statistical Analysis: Wilcoxon matched-pairs signed rank test, ****<0.0001. k. Analysis of percentage of cells expressing ADAR2 (I) with or without TTX. N=3 independent dissections and n=15 fields of view. Statistical Analysis: Unpaired t-test, *<0.05.

### Complete and partial ADAR2 knockdown differentially alter GluK2 and GluA2 Q/R editing

To further examine the role of ADAR2 in KAR scaling we generated two different shRNAs against ADAR2. One shRNA we named shRNA_‘Complete’_ ablated essentially all ADAR2 whereas another shRNA that we called shRNA_‘Partial’_, reduced ADAR2 levels to ~50% of the control (Figure 3A,B), comparable to the loss observed with TTX. Furthermore, shRNA_‘Partial’_ knockdown reduced the intensity of signal and percentage of cells expressing ADAR2 to levels similar to those elicited by TTX treatment (Figure 3C,D,E).

**Figure 3:**
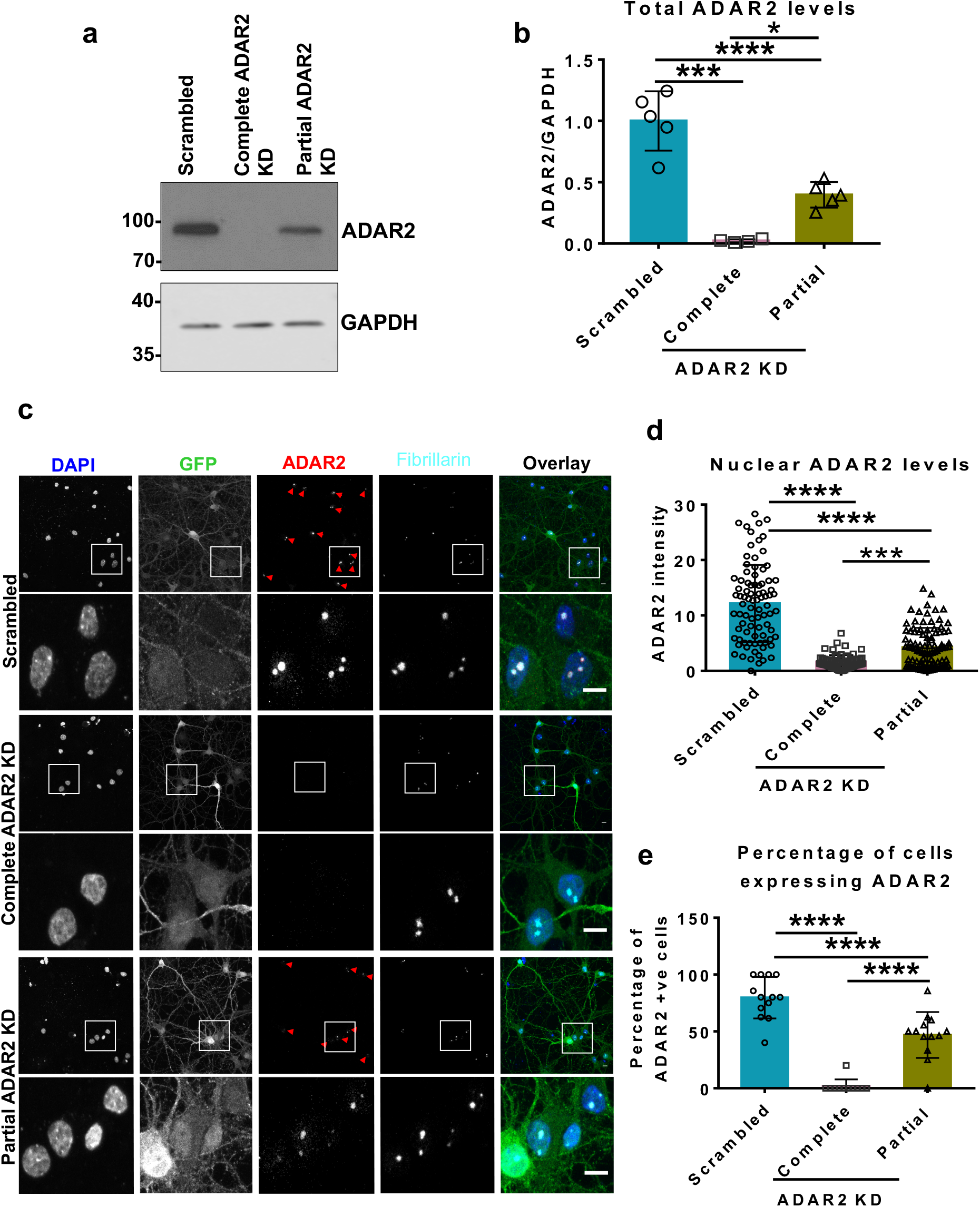
Complete and partial ADAR2 knockdown. a. Representative western blots of total ADAR2 and GAPDH levels in hippocampal neurons treated with either scrambled, complete or partial ADAR2 KDs. b. Quantification of (A) total ADAR2 normalised to GAPDH from 5 independent experiments for scrambled and partial knockdown and 4 independent experiments for complete knockdown. Statistical Analysis: One-Way ANOVA with Tukey’s multiple comparisons test; *<0.05, ***<0.001, ****<0.0001. c. Representative images of hippocampal neurons imaged for DAPI (blue), GFP (green, lentivirus infected cells), ADAR2 (red) and Fibrillarin (nucleolar marker; cyan) for cells infected with either scrambled, complete or partial ADAR2 KD. Bottom panels show zoom in images as indicated and the red arrows indicate cells expressing ADAR2. d. Quantification of (C) ADAR2 intensity per nucleus. N=3 independent dissections and n= 75 cells (complete KD), 86 cells (scrambled) and 99 cells (Partial knockdown). Statistical Analysis: One Way Anova with Tukey’s multiple comparisons test, ***<0.001, ****<0.0001. e. Quantification of (C) percentage of cells expressing ADAR2. N=3 independent dissections and n=11-13 fields of view. Statistical Analysis: One Way Anova with Tukey’s multiple comparisons test, ****<0.0001.

We next compared how ADAR2 knockdown affected GluK2 and GluA2 editing. As expected, shRNA_‘Complete’_ ADAR2 knockdown reduced GluK2 Q/R editing by over 60%, whereas shRNA_‘Partial’_ knockdown of ADAR2 only reduced GluK2 Q/R editing by ~20% (Figure 4A,B). DNA sequencing chromatographs from cDNA of cells treated with shRNA_‘Complete’_ show a dramatic change in the base read of the editing site to CAG (Q) rather than CGG (R), whereas neurons treated with shRNAPartar show a mixture of both CGG and CAG (Figure 4C). Interestingly, shRNA_‘Partial’_ knockdown of ADAR2 had no effect on the Q/R editing of the AMPAR subunit GluA2 while the shRNA_‘Complete’_ knockdown only reduced GluA2 editing by ~30% (Figure 4D,E,F). These data indicate that GluK2 editing levels are uniquely sensitive to ADAR2 changes and support a model whereby loss of ADAR2 during homeostatic scaling directly promotes surface expression of GluK2 containing KARs through a reduction in GluK2 editing.

**Figure 4:**
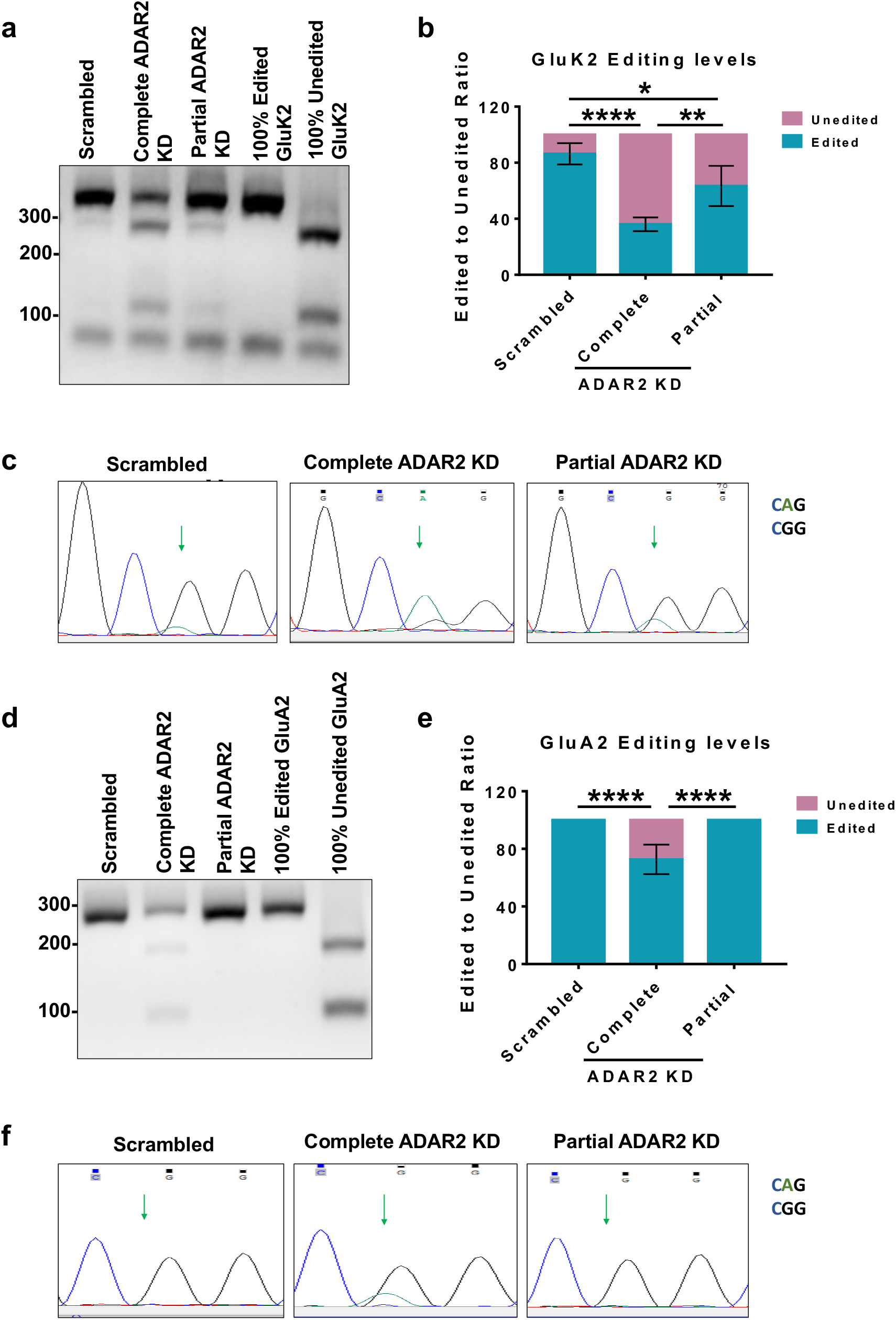
Complete and partial ADAR2 knockdown differentially alter GluK2 and GluA2 Q/R editing. a. RT-PCR and BbvI digestion analysis of GluK2 Q/R editing from hippocampal neurons infected with either scrambled, complete or partial ADAR2 KDs. b. Quantification of (A) from 4 independent experiments. Statistical Analysis: One Way ANOVA with Tukey’s multiple comparisons test; *<0.05, **<0.01, ****<0.0001. c. Sanger sequencing chromatographs of the GluK2 PCR products from hippocampal neurons infected with either scrambled, complete or partial ADAR2 KDs, showing dual A and G peaks at the editing site indicated by the green arrows. The green peak represents an A base read and black represents a G base read. d. RT-PCR and BbvI digestion analysis of GluA2 Q/R editing from hippocampal neurons infected with either scrambled, complete or partial ADAR2 KDs. e. Quantification of (D) from 4 independent experiments. Statistical Analysis: One Way ANOVA with Tukey’s multiple comparisons test; ****<0.0001. f. Sanger sequencing chromatographs of the GluA2 PCR products from hippocampal neurons infected with either scrambled, complete or partial ADAR2 KDs. Green arrows indicate the editing site. Only samples treated with complete ADAR2 knockdown show a dual A and G peak at the editing site. The green peak represents an A base read and black represents a G base read.

### Partial ADAR2 knockdown phenocopies and occludes TTX upscaling of GluK2 containing KARs

Intriguingly, shRNA_‘Partial’_ reduces ADAR2 to levels comparable to those following TTX treatment and results in a similar shift in the proportions of edited to unedited GluK2. We therefore wondered if shRNA_‘Partial’_ alone is sufficient to mediate KAR upscaling. Partial ADAR2 knockdown significantly increased GluK2 surface expression with no effect on EGFR surface expression (Figure 5A,B,C), indicating that reduced levels of ADAR2 result in upscaling GluK2 containing KARs in the absence of TTX treatment. TTX treatment in combination with shRNA_‘Partial’_ knockdown was not additive (Figure 5A,B) consistent with TTX induced upscaling being mediated by a reduction in ADAR2 levels. In shRNA_‘Partial’_ ADAR2 knockdown neurons, 24 h TTX treatment did elicit a further decrease in ADAR2 levels (SI Figure 2A.B), indicating that even when already depleted, ADAR2 levels are still subject to activity-dependent regulation. However, despite the summative decrease in ADAR2 levels, shRNA_‘Partial’_ ADAR2 knockdown combined with TTX treatment did not further decrease GluK2 Q/R editing compared to either treatment alone (Figure 5D,E). These data indicate that loss of ADAR2 is sufficient to mediate TTX induced up-scaling of KARs.

**Figure 5:**
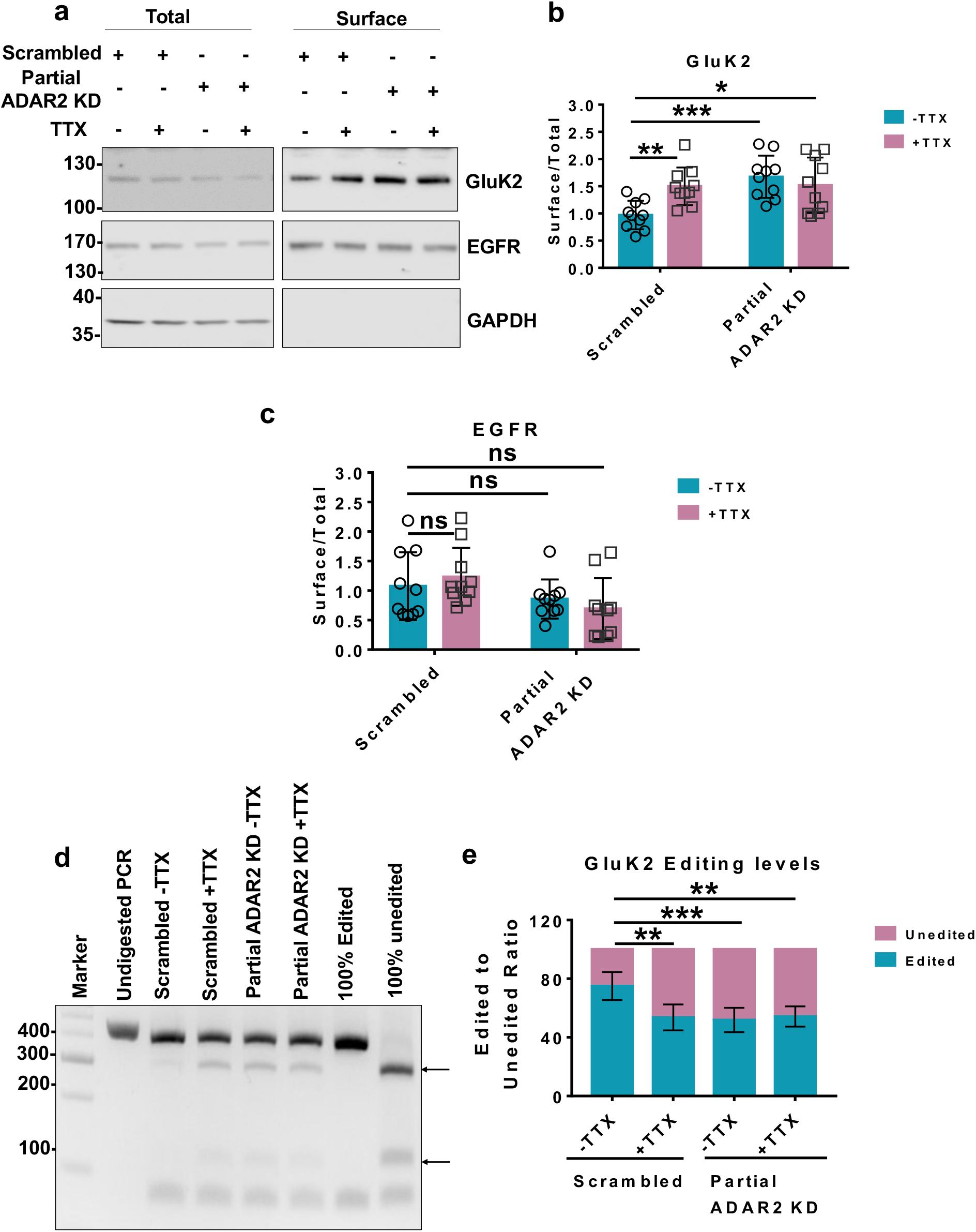
Partial ADAR2 knockdown phenocopies and occludes TTX up-scaling of GluK2 containing KARs. a. Representative western blot of total and surface levels of GluK2, EGFR and GAPDH in scrambled or ADAR2 KD infected cells in the presence or absence of TTX. EGFR was used as a negative control while GAPDH was used as a control to show only surface proteins were labelled with biotin. b. Quantification of (A) surface levels of GluK2 from 10 independent experiments. Surface levels were normalised to their total levels. Statistical Analysis: Two-Way ANOVA with Tukey’s Multiple comparisons test; *<0.05, **<0.01, ***<0.001. c. Quantification of (A) surface levels of EGFR from 10 independent experiments. Surface levels were normalised to their total levels. Statistical Analysis: Two-Way ANOVA with Tukey’s Multiple comparisons test; ns>0.05. d. RT-PCR and BbvI digestion analysis of GluK2 Q/R editing from hippocampal neurons infected with either scrambled or partial ADAR2 KD with or without TTX. e. Quantification of (D) from 5 independent experiments. Statistical Analysis: One Way ANOVA with Tukey’s multiple comparisons test; **<0.01, ***0.001.

### Loss of ADAR2 during scaling is not dependent on its interaction with Pin1

The nuclear protein Pin1 retains ADAR2 in the nucleus and interfering with this allows ADAR2 export to the cytosol, its ubiquitination and degradation (32). Our RT-qPCR experiments show that total ADAR2 mRNA levels are unaffected by TTX, indicating that scaling is not transcriptionally mediated (SI Figure 3). We therefore wondered if destabilising the Pin1-ADAR2 protein interaction underpins ADAR2 loss by TTX but Pin 1 levels were unchanged following TTX treatment (Figure 6A,B).

**Figure 6:**
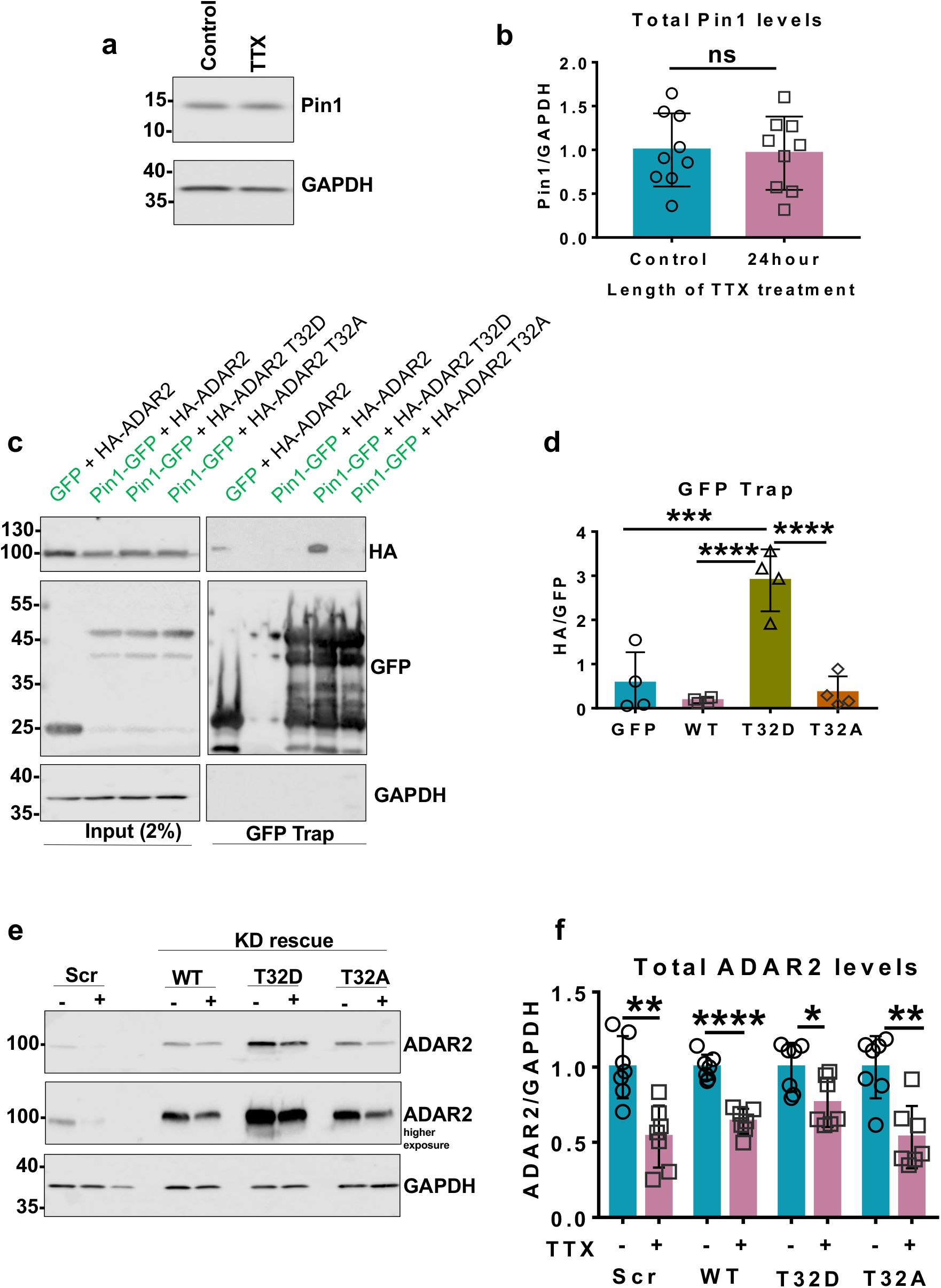
Loss of ADAR2 during TTX mediated scaling is not dependent on Pin1 or ADAR2 phosphorylation. a. Representative western blots of total Pin1 and GAPDH levels in neurons with or without TTX treatment for 24h. b. Quantification of (A) Pin1 levels normalised to GAPDH from 9 independent experiments. Statistical Analysis; Unpaired t-test: ns>0.05. c. Representative blots of GFP-trap performed in HEK293T cells where Pin-GFP was overexpressed with either HA-WT ADAR2, HA-T32D ADAR2 or HA-T32A ADAR2. The pulldowns were blotted for HA, GFP and GAPDH. Free GFP was overexpressed with WT-ADAR2 as a negative control. d. Quantification of (C) showing interaction of HA-T32D ADAR2 with Pin1-GFP. The HA signal was normalised to the GFP. N=4 independent experiments. Statistical Analysis: One-way ANOVA with Tukey’s multiple comparisons test; ***<0.001, ****<0.0001. e. Representative western blots of total ADAR2 and GAPDH in cells infected with either scrambled or KD rescue lentiviruses expressing either WT ADAR2, T32D ADAR2 or T32A ADAR2, in the presence or absence of TTX. f. Quantification of (E) total ADAR2 levels normalised to GAPDH from 7 independent experiments. Each TTX treated condition was normalised to their respective non-treated control. Statistical Analysis: Unpaired t-test; *<0.05, **<0.01, ***<0.00, ****<0.0001.

We next investigated the role of ADAR2 phosphorylation at threonine 32 (T32) in regulating ADAR2 levels during scaling, since this has been reported to crucial for the ADAR2-Pin1 interaction (32). In contrast to WT and phosphonull (T32A) ADAR2, the phosphomimetic (T32D) ADAR2 mutant binds very strongly to Pin1 in GFP-trap assays (Figure 6C,D). We therefore tested if the phosphonull or phosphomimetic ADAR2 mutants were more sensitive to TTX treatment. We first knocked down endogenous ADAR2 and replaced it with HA-tagged WT ADAR2 (SI Figure 4A,B). More than 80% of the cells expressed the knockdown-rescue, similar to the percentage of untreated neurons that express ADAR2 (SI Figure 4A,B). We then investigated the stability of the phosphonull or phosphomimetic ADAR2 mutants in response to TTX treatment. Similar to WT ADAR2, levels of both mutants were significantly decreased by TTX treatment (Figure 6E,F; SI Figure 5). Furthermore, nuclear levels of all the constructs decreased following TTX treatment (SI Figure 6A,B). Thus, activity-dependent untethering of ADAR2 from Pin1 is not a primary mechanism responsible for the loss of ADAR2 during TTX mediated upscaling.

### TTX mediated scaling enhances proteasomal degradation of ADAR2

It has been proposed that ADAR2 is ubiquitinated and degraded in the cytosol (32). We therefore tested the effects of TTX on ADAR2 stability in the presence or absence of the proteasomal inhibitor Bortezomib (BTZ) (33). Consistent with suppression of synaptic activity leading to enhanced proteasomal degradation, BTZ prevented TTX-evoked decreases in ADAR2 (Figure 7A,B) and resulted in the accumulation of ubiquitinated products (Figure 7A,C). We next performed nuclear and cytoplasmic fractionations to determine if ADAR2 is exported from the nucleus for degradation in the cytosol. BTZ prevented the TTX-evoked decrease in ADAR2 in both the nuclear and cytosolic fractions, and actually led to a significant accumulation of ADAR2 in the cytosol (Figure 7D,E,F). Thus, ADAR2 may be exported to the cytosol to be ubiquitinated or ubiquitinated in the nucleus and exported to the cytosol to be degraded. In either case TTX induces proteasomal degradation of ADAR2.

**Figure 7:**
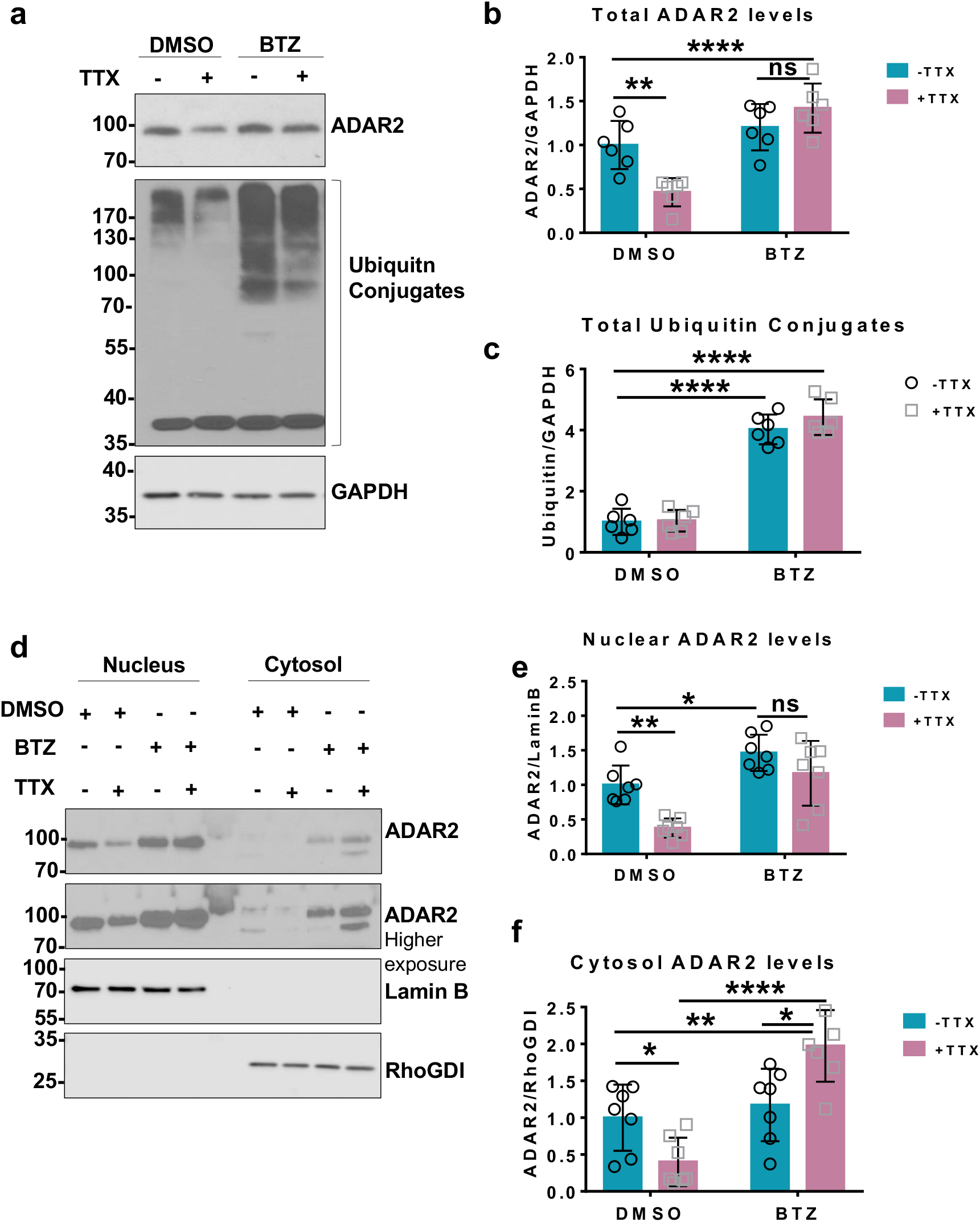
TTX mediated scaling enhances proteasomal degradation of ADAR2. a. Representative western blots of total ADAR2, total ubiquitin and GAPDH levels in neurons treated with either DMSO or 1μM Bortezomib (BTZ) for 20 h either in the presence or absence of 24 h TTX. b. Quantification of (A) total ADAR2 immunoblots normalised to GAPDH from 6 independent experiments. Statistical Analysis: Two-way ANOVA with Tukey’s multiple comparisons test: **<0.01, ****<0.0001, ns>0.05. c. Quantification of (A) total ubiquitin conjugated products normalised to GAPDH from 6 independent experiments. Statistical Analysis: Two-way ANOVA with Tukey’s multiple comparisons test: ****<0.0001. d. Representative western blots of ADAR2 in the nucleus and cytoplasm in the presence of DMSO or BTZ with or without TTX treatment. Lamin B was used as a nuclear marker and RhoGDI as cytosol marker. e. Quantification of (D) nuclear ADAR2 from 7 independent experiments. Nuclear ADAR2 levels were normalised to Lamin B. Statistical Analysis: Two-way ANOVA Tukey’s multiple comparisons test: *<0.05, **<0.01, ns>0.05. f. Quantification of (D) cytosolic ADAR2 from 7 independent experiments. Cytosolic ADAR2 was normalised to RhoGDI level. Statistical Analysis: Two-way ANOVA with Tukey’s multiple comparisons test: *<0.05, **<0.01, ****<0.0001.

### Blocking ADAR2 degradation prevents KAR up-scaling in response to TTX

Since BTZ prevents the loss of ADAR2 in the nucleus in response to TTX, we next tested if preventing ADAR2 degradation with BTZ blocks TTX up-scaling of GluK2. Indeed, surface biotinylation showed that BTZ prevents TTX-induced GluK2 upscaling (Figure 8A,B) with no effect on EGFR (Figure 8A,C). These results support the hypothesis that ADAR2 degradation mediates KAR up-scaling.

**Figure 8:**
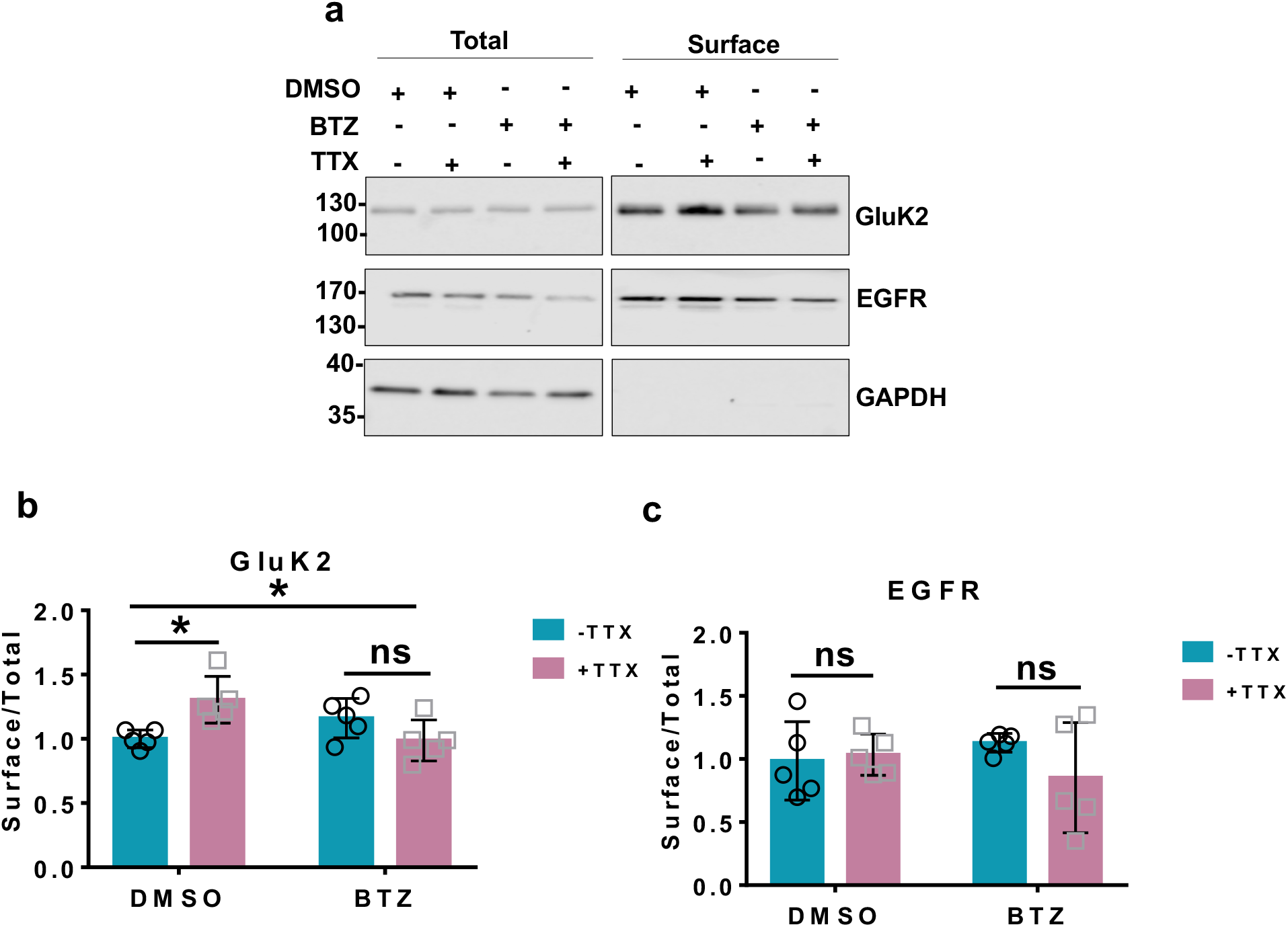
Blocking ADAR2 degradation with BTZ prevents KAR up-scaling in response to TTX. a. Representative western blots of total and surface levels of GluK2, EGFR and GAPDH in DMSO and BTZ (20 h, 1μM) treated cells in the presence or absence of 24 h TTX. EGFR was used as a negative control while GAPDH was used as a control to show only surface proteins were labelled with biotin. b. Quantification of (A) surface levels of GluK2 from 5 independent experiments. Surface levels were normalised to their total levels. Statistical Analysis: Two-Way ANOVA with Tukey’s Multiple comparisons test; *<0.05, ns>0.05. c. Quantification of (A) surface levels of EGFR from 5 independent experiments. Surface levels were normalised to their total levels. Statistical Analysis: Two-Way ANOVA with Tukey’s Multiple comparisons test; ns>0.05.

## Discussion

### Selective ADAR2 mediated Q/R editing of GluK2 containing KARs regulates up-scaling

Similar to AMPARs and NMDARs, GluK2 containing KARs up-scale following chronic suppression in synaptic activity (25). Here we show that the same TTX treatment leads to proteasomal degradation of ADAR2 and, consequently, reduced Q/R editing of GluK2. These data are consistent with activity-dependent loss of ADAR2 leading to the alteration in KAR editing which, in turn, directly mediates KAR up-scaling via increased KAR assembly and ER exit of unedited GluK2(Q) compared to edited GluK2(R) (9, 25).

Partial ADAR2 knockdown mimicked the decrease in ADAR2 levels observed after TTX scaling and did not alter GluA2 editing. We therefore used this tool to further study the role of ADAR2 in GluK2 up-scaling. Partial loss of ADAR2 phenocopies and occludes TTX-mediated GluK2 editing and upscaling. Thus, loss of ADAR2 is sufficient to up-scale KARs in the absence of TTX, and strongly suggests that reduced KAR editing underpins KAR up-scaling (Figure 9 schematic).

**Figure 9:**
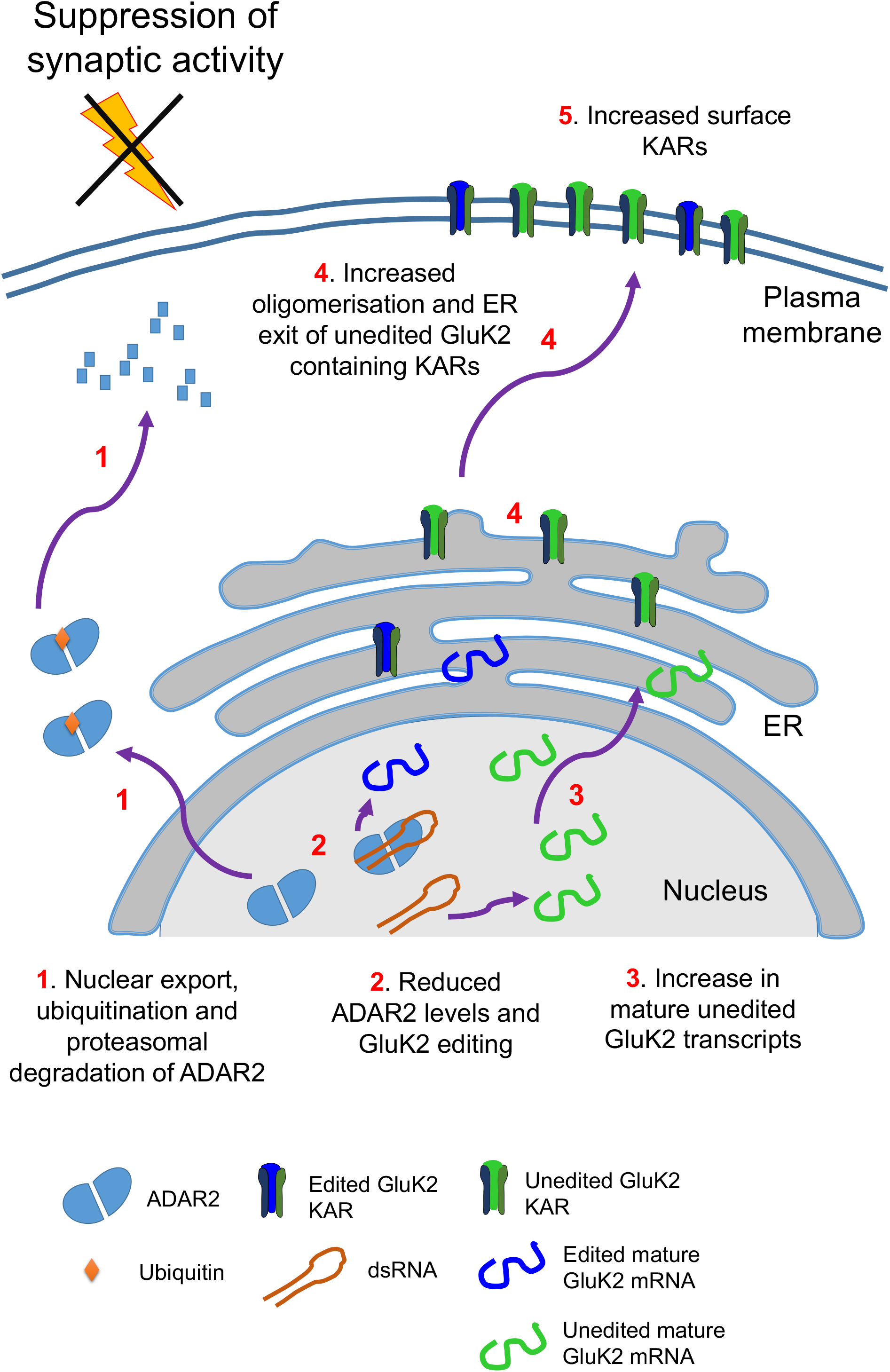
Schematic of ADAR2 mediated Q/R editing regulating GluK2 containing KARs homeostatic up-scaling. Under basal conditions unedited GluK2 transcripts are edited at their Q/R site by ADAR2 enzyme resulting in ~80% edited mature GluK2 transcripts. The resultant edited and unedited GluK2 subunits oligomerise in the ER and traffic into the surface. Under conditions of synaptic activity suppression with TTX treatment, ADAR2 undergoes proteasomal degradation in the cytosol (1). This results in less ADAR2 editing of GluK2 pre-mRNA transcripts (2) and increased levels of unedited GluK2 mature transcripts (3). The subsequent increase in the proportion of unedited GluK2(Q) allows enhanced oligomerisation and ER exit (4) to increase surface expression of GluK2 containing KARs on the surface (5).

### Differential sensitivity of GluK2 to changes in ADAR2 levels

While we observed robust up-scaling of both KARs (GluK2) and AMPARs (GluA2) by TTX, the mechanisms are different because the Q/R editing status of GluA2 was not altered. In line with this, complete loss of ADAR2 decreased GluA2 editing by ~30% and GluK2 editing by >60%, whereas partial ADAR2 loss decreased editing of GluK2 by ~20% but had no discernible impact on GluA2 editing. Notably, it has been reported that ADAR2 regulation of GluA2 is specifically confined to CA1 hippocampal and does not occur in CA3 (34). GluA2 is preferentially and almost completely Q/R edited, even when only low levels of ADAR2 are available. GluK2 mRNA, on the other hand, is more sensitive to changes in the levels of ADAR2. We interpret these data to indicate that the Q/R editing of KARs is more dynamically and activity-dependently regulated than AMPARs, and reason that this provides a rapidly tunable system to control KAR forward trafficking and scaling.

### Activity dependent regulation of ADAR2 proteostasis during TTX treatment

ADAR2 acts in the nucleus where it is stabilised by Pin1. Loss of the ADAR2-Pin1 interaction leads to ADAR2 export into cytosol where it is ubiquitinated and degraded (32). Pin1 mediated stabilisation is important for ADAR2 editing activity during development in cortical neurons (11). However, our experiments suggest Pin1-ADAR2 interactions do not play a role in TTX up-scaling since both phosphonull and phosphomimetic mutants of ADAR2, which decrease or enhance binding to Pin1 respectively, were equally susceptible to the TTX mediated loss.

TTX induces ADAR2 degradation which is blocked by the proteasome inhibitor BTZ. We speculate that ADAR2 accumulated in the cytosol can be re-imported into nucleus to maintain a functional pool of nuclear ADAR2. Consistent with this, blocking proteasomal degradation of ADAR2 prevents TTX induced GluK2 up-scaling. Thus, the loss of ADAR2 and consequent reduction in GluK2 Q/R editing are necessary and sufficient for KARs up-scaling.

KARs play key roles in neuronal excitability, synaptic transmission and plasticity, and network activity (3, 35). Moreover, KAR dysregulation is implicated in neuronal pathologies (36, 37). How KAR upscaling impacts on these pathways is an outstanding question. Our data provide novel mechanistic insight into how neuronal activity controls the surface expression and properties of KARs during homeostatic scaling that will have wide implication for neuronal function, and dysfunction in disease.

## Materials and Methods

### Primary neuronal cultures

Primary rat hippocampal neurones were dissected from E18 Wistar rat pups as previously described (38). Briefly neurones were dissected from E18 Wistar rats followed by trypsin dissociation and cultured for up to 2 weeks. For the first 24 h, cells were grown in plating media: Neurobasal media (Gibco) supplemented with 10% horse serum (Sigma), B27 (1x, Gibco), P/S (100 units penicillin and 0.1mg/ml streptomycin; ThermoScientific) and 5mM Glutamax (Gibco). After 24 h, plating media was replaced with feeding media (Neurobasal containing 2mM Glutamax and lacking horse serum and P/S). For biochemistry experiments, cells were plated at a density of 500,000 per 35mm well and 250,000 per coverslip for imaging experiments.

### ADAR2 cloning

ADAR2 was cloned from rat neuronal cDNA and ADAR2 shRNA knockdown and knockdown-rescue viruses were generated as previously described (38). ADAR2 was cloned from rat neuronal cDNA into the KpnI and XbaI sites of the vector pcDNA3 with a HA tag at its N-terminus. Phosphomutants of ADAR2 were generated by site-directed mutagenesis. Pin1 was cloned from rat neuronal cDNA into the EcoRI and BamHI sites of the vector pEGFP-N1.

### Lentivirus generation

For ADAR2 knockdown experiments, shRNA sequences targeting ADAR2 cloned into a modified pXLG3-GFP vector (38) under the control of a H1 promoter. The ADAR2 target sequences were:

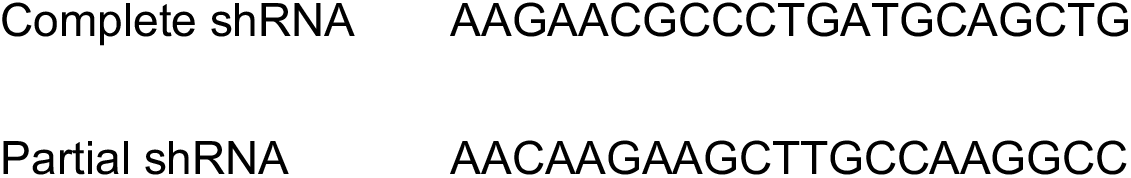

For rescue experiments, shRNA-insensitive HA-ADAR2 was cloned into a modified pXLG3-GFP vector under the control of an SFFV promoter.

The viruses were produced in HEK293T cells as reported previously (38), harvested and added to DIV 9/10 hippocampal neurons for 5 days and lysed accordingly. For 24 h treatment experiments, cells were treated with on the 4^th^ day after virus addition and harvested accordingly on the 5^th^ day after the completion of the time course.

### Scaling, developmental and TTX timecourse and BTZ treatment

For scaling experiments, cells were treated with 1μM TTX (Tocris) for 24 h and were either lysed directly in 1x sample buffer and heated for 10 minutes at 95°C or were used for either surface biotinylation or fractionation experiments (see below). For the TTX timecourse, cells were harvested directly into 1x sample buffer (4 x sample buffer (0.24M Tris-HCL, 8% SDS, 40% glycerol, 10% β-mercaptoethanol and 0.009% Bromophenol blue) diluted in water). In experiments inhibiting proteosomal degradation, cells were treated with bortezomib (BTZ; Cell Signalling) dissolved in DMSO for 20 h at 1 μM concentration. The control cells were treated with an equal volume of DMSO.

### Cell surface biotinylation and streptavidin pulldown

Cell surface biotinylation was performed essentially as previously described (25). All steps were performed on ice with ice-cold buffers unless stated otherwise. Live hippocampal neurones post stated treatments were washed twice in phosphate buffered saline (PBS). Surface proteins were labelled with membrane impermeable Sulfo-NHS-SS biotin (0.3mg/ml, Thermo Scientific) for 10 mins on ice and washed 3x with PBS. 100mM NH_4_Cl was added to quench free biotin-reactive groups and cells were extracted with lysis buffer (50mM Tris pH 7.4, 150mM NaCl, 1% triton, 0.1% SDS, protease inhibitors (Roche)), incubated on ice for 30 mins and centrifuged (15,000g, 4°C, 20 mins) to remove cell debris. For streptavidin pulldown, each lysate was added to 30μl of streptavidin beads (Sigma) and left on a wheel to rotate for 90 minutes at 4°C. The beads were then washed 3x with wash buffer (lysis buffer without protease inhibitors) and proteins eluted with 2x sample buffer and boiled for 10 minutes at 95°C. The samples were then resolved by SDS-PAGE and immunoblotted.

### Subcellular fractionation

All the steps were performed on ice with ice-cold buffers unless stated otherwise. Following stated treatments, the cells were washed with PBS followed by addition of buffer 1 (150mM NaCl, 50mM HEPES pH 7.4, 25μg/ml Digitonin and protease inhibitors), incubated for 20 minutes, scraped, homogenised and centrifuged at 15000g for 30 minutes at 4°C. The supernatant consisted of the cytosolic proteins, while the pellet was resuspended in buffer 2 (Buffer 1 with 1% triton), incubated for 20 minutes and again centrifuged at 15000g for 30 minutes at 4°C. The supernatant consisted of mitochondrial proteins while the pellet was resuspended in buffer 3 (150mM NaCl, 50mM HEPES pH 7.4, 0.5% sodium deoxycholate, 0.1% SDS, protease inhibitors and 0.5% triton), incubated for 1 h and centrifuged at 15000g for 30 minutes at 4°C. The supernatant was discarded and the pellet, consisting of nuclear proteins, was resuspended in buffer 3. The cytosolic supernatant was concentrated using 4 volumes of acetone (kept at −20°C), incubated at −20°C for 1 h and spun for 20 minutes at 1500g and resuspended in buffer 3. BCA assay was then performed to determine protein concentrations and allow equal loading.

### BCA Assay

BCA Assay was performed using a commercial kit (Pierce, Thermo Scientific). Following 30 minutes incubation at 37°C, the samples were read using a plate reader (Versamax Microplate reader, Molecular devices) at a wavelength of 562nm.

### Western blotting and antibodies

Antibodies used: GluK2 (1:1000, Millipore, rabbit polyclonal), GluA2 (1:1000, Synaptic Systems, rabbit polyclonal), ADAR2 (1:1000, Sigma, rabbit polyclonal), GAPDH (1:10,000, Abcam, mouse monoclonal), RhoGDI (1:1000, Abcam, rabbit polyclonal), Lamin B (1:1000, Santa Cruz, Goat polyclonal), EGFR (1:1000, Abcam, rabbit polyclonal), HA (1:2000, mouse, Sigma), Pin1 (mouse monoclonal, 1:1000, Santa cruz), ADAR1 (mouse monoclonal, 1;1000, Santa Cruz) and GFP (rat monoclonal, 1:10,000, Chromotek). Western blots were imaged and quantified using LI-COR Image Studio software or developed on X-ray film in a dark room using developer and fixer solutions. The blots were then scanned and quantified using FIJI ImageJ studio. Surface levels were normalised to their respective total levels. Nuclear protein levels were normalised to LaminB and cytosolic protein levels to RhoGDI. Treated samples were normalised to their control samples.

### RNA extraction, RT-PCR, and BbvI digestion and RT-qPCR

RNA samples were extracted from DIV14/15 hippocampal neurons following the stated treatments using RNeasy Mini Kit (Qiagen) according to the manufacturer’s instructions. 1 μg of RNA was used per condition and reverse transcribed to cDNA using RevertAid First Strand cDNA Synthesis Kit (Thermo Scientific) following the manufacturer’s instructions. The following primers (spanning the M2 region of GluK2(39) and GluA2) were used, giving a PCR product of 452bp and 252bp:

> GluK2 F: 5’-GGTATAACCCACACCCTTGCAACC-3’
>
> GluK2 R: 5’-TGACTCCATTAAGAAAGCATAATCCGA-3’
>
> GluA2 F: 5’-GTGTTTGCCTACATTGGGGTC-3’
>
> GluA2 R: 5’-TCCTCCTACACGG CTAACTTA-3’

5μl of cDNA was used to set up PCR reactions [50μl total, 35 cycles, 20s denaturing at 95°C, 10s annealing at 60°C and 15 s elongation at 70°C].

To determine the level of RNA editing, BbvI (New England Biolabs) digestion was used as previously (39). Total 20μl digestion was set up using 10μl of PCR product at 37°C for 2 h. All of the digested product was run on 4% agarose gel and the ethidium bromide stained bands were imaged using UV transilluminator and quantified using FIJI NIH ImageJ. To determine the level of editing in GluK2, the following formula was used: [Intensity of 376 (edited)/Intensity of (376 (edited) + 269 (unedited))]*100. The band at 76bp allowed to determine equal loading. For GluA2, [Intensity of 158 (edited)/Intensity of (158 (edited) + 94 (unedited))]*100.

Purified PCR products were also sent for sequencing to Eurofins Genomics at 4ng/μl along with the above GluK2 and GluA2 F primers, to obtain sequence chromatographs.

For RT-qPCR, 2μl of the cDNA samples per condition were mixed with PowerUp SYBR green Master Mix (Life Technologies) and forward and reverse primers targeting ADAR2 and GAPDH and amplified quantitatively using Real Time PCR System (MiniOpticon, BioRAD) for 40 cycles and Ct values were recorded. Each reaction was performed in triplicate and average Ct was measured per condition. ADAR2 Ct values were normalised to GAPDH Ct values and ADAR2 mRNA fold difference value of TTX treated conditions was normalised against the untreated control. Melting curve of the primers were also determined to ensure the specificity of the primers and lack of primer dimer formation. The primers used were: ADAR2 F – TCCCGCCTGTGTAAGCAC, GAPDH AAT CCCAT CACCAT CTT CCA.

### GFP trap

GFP-trap protocols were as previously published (40). HEK293T cells were transfected the next day using Lipofectamine™3000 and 2.5μg of each construct. 48 h post-transfection, cells were washed with PBS and lysed and harvested in lysis buffer (20mM Tris pH7.4, 137mM NaCl, 2mM sodium pyrophosphate, 2mM EDTA, 1% triton X-100, 0.1% SDS, 25mM β-glycerophosphate, 10% glycerol, protease inhibitors (Roche), phosphatase inhibitor cocktail 2 (1:100, Sigma)). The lysates were left to incubate for 30 minutes on ice and centrifuged at 1500g for 20 minutes at 4°C to remove any cell debris. The supernatant was then added to 5μl of GFP-trap beads (Chromotek), incubated on wheel at 4°C for 90 minutes and washed 3x with wash buffer (lysis buffer without protease or phosphatase inhibitors). The samples were then lysed in 2x sample buffer, heated at 95°C for 10 minutes and separated using SDS-PAGE.

### Fixed immunostaining, imaging and analysis

Immunostaining was performed as previously described (41). For fixed immunostaining, cells post TTX treatment or lentiviral treatment as indicated were fixed with 4% formaldehyde for 10 minutes, washed 3 times with PBS, treated with 100mM glycine to quench any remaining formaldehyde and washed 3 times with PBS. The cells were permeabilised and blocked with 3% BSA in PBS and 0.1% triton for 30 minutes. The cells were then incubated for 1 h with primary antibodies (anti-ADAR2 (Abcam, rabbit) 1:400, anti-Fibrillarin (Abcam, mouse) 1:400 and anti-HA (Sigma, mouse) 1:600) in 3% BSA at room temperature, washed 3 times for 5 minutes each with PBS and incubated for 45 minutes with the indicated secondary antibodies (Jackson Immunoresearch Antibodies, 1:400) in 3% BSA at room temperature. Three x 5 minute washes were performed with PBS and the cells were mounted using DAPI containing fluoromount.

A Leica SP7 confocal microscope was used to image the coverslips. FIJI NIH Image J was used to compress the z-stacks and analyse the mean intensity per nucleus using the DAPI channel to draw regions of interest. To calculate the percentage of cells expressing ADAR2, all the cells expressing ADAR2 were manually counted per image taken.

### Statistical Analysis

Mean value were calculated for all data and all error bars show standard deviation. All statistical analysis was performed using GraphPad Prism software version 7.0 as stated.

## Acknowledgements

We are grateful to the MRC, BBSRC and Wellcome Trust for financial support. SG and AJE were supported by Wellcome Trust Dynamic Cell Biology PhD studentships. We thank Dr Yasuko Nakamura for excellent technical and logistical support.

## Author Contributions

SG performed all of the experiments. AJE and KAW provided specialised constructs, advice and assisted in some experiments. JMH supervised the research. All authors contributed to writing the manuscript.

---

**Supplementary Figure 1:**
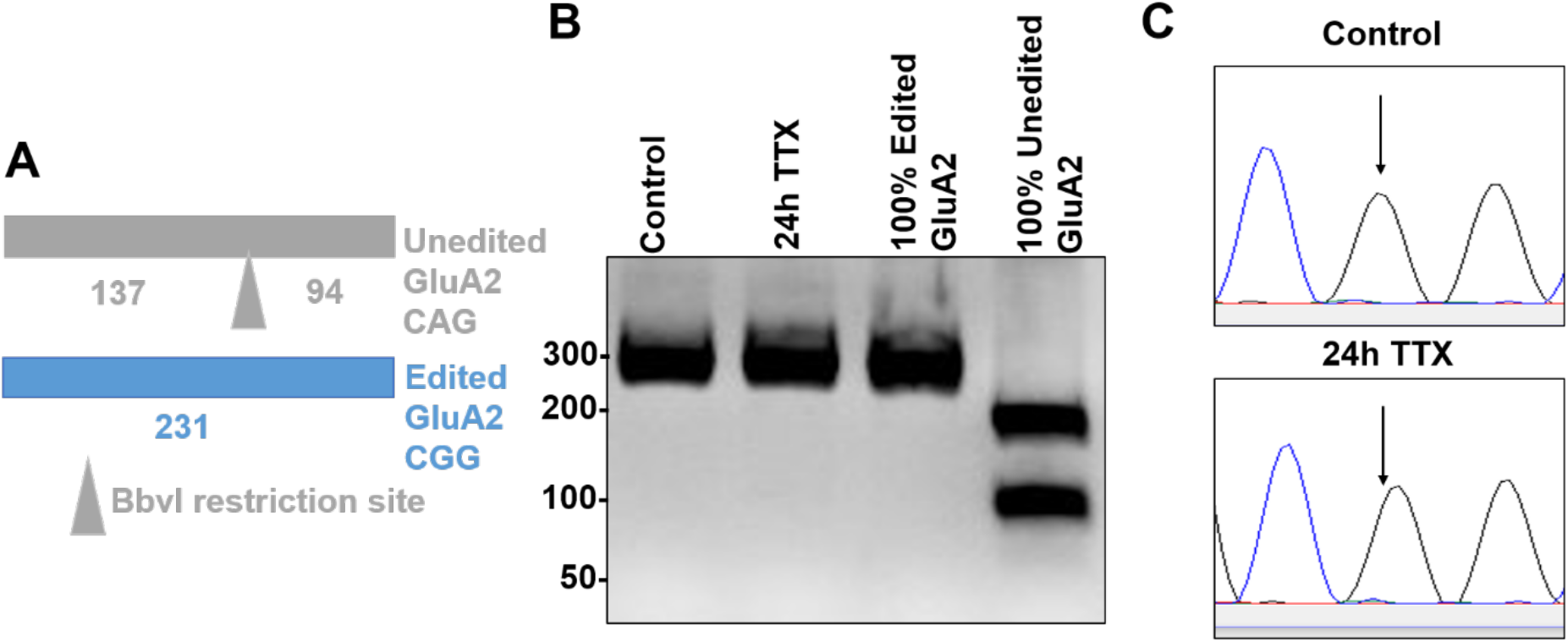
GluA2 editing status does not change with TTX treatment despite the loss of ADAR2 levels. A. Schematic of BbvI digestion analysis on the PCR amplified M2 region of GluA2. B. RT-PCR and BbvI digestion analysis of GluA2 Q/R editing from hippocampal neurones treated with or without TTX. C. Sanger sequencing chromatographs of the GluA2 PCR products from hippocampal neurones treated with or without TTX, showing no changes in the editing levels. Black arrows indicate the editing site. Green peak represents A base read and black represents G base read.

**Supplementary Figure 2.**
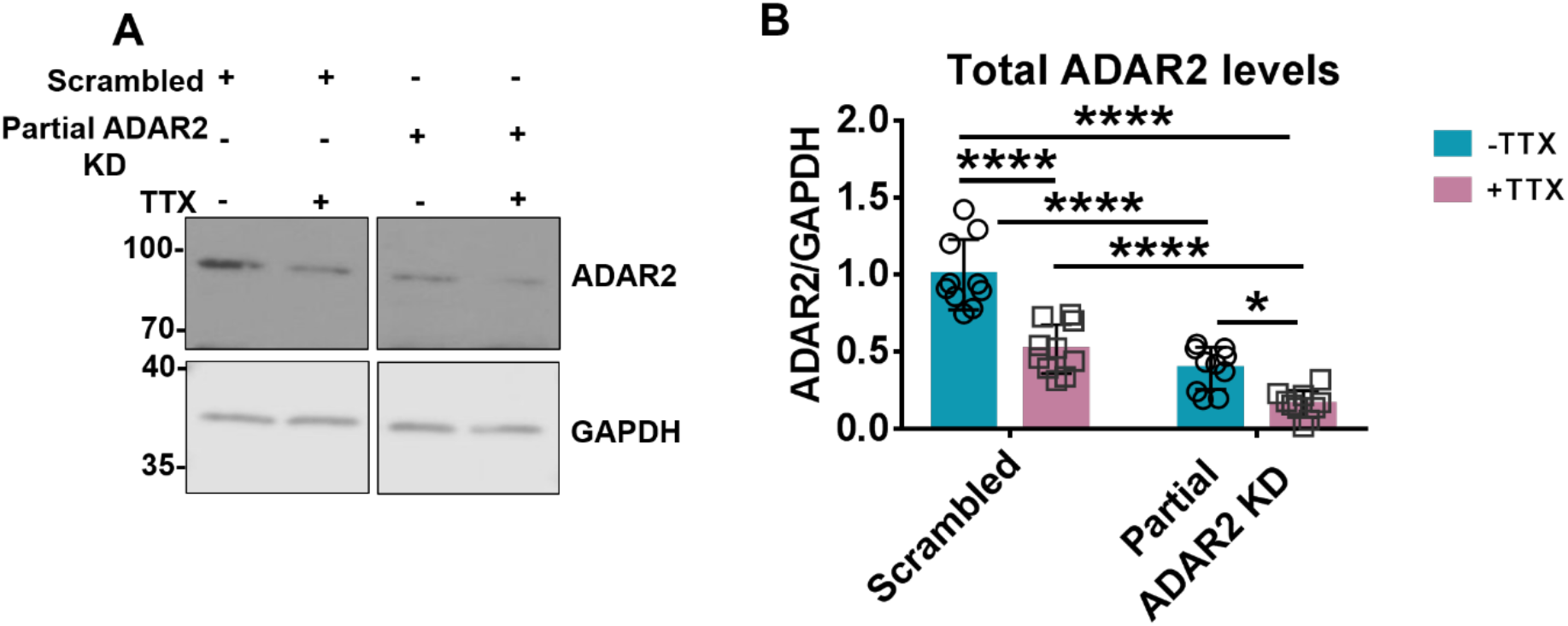
ADAR2 level further decreases with TTX treatment in an already depleted system. A. Representative western blots of total ADAR2 and GAPDH levels in cells infected with either scrambled or Partial ADAR2 KD either in the presence or absence of 24h TTX. B. Quantification of the (A) total ADAR2 immunoblots normalised to GAPDH from 10 independent experiments. Statistical Analysis: Two-way ANOVA with Tukey’s multiple comparisons test: *<0.05, ****<0.0001.

**Supplementary Figure 3.**
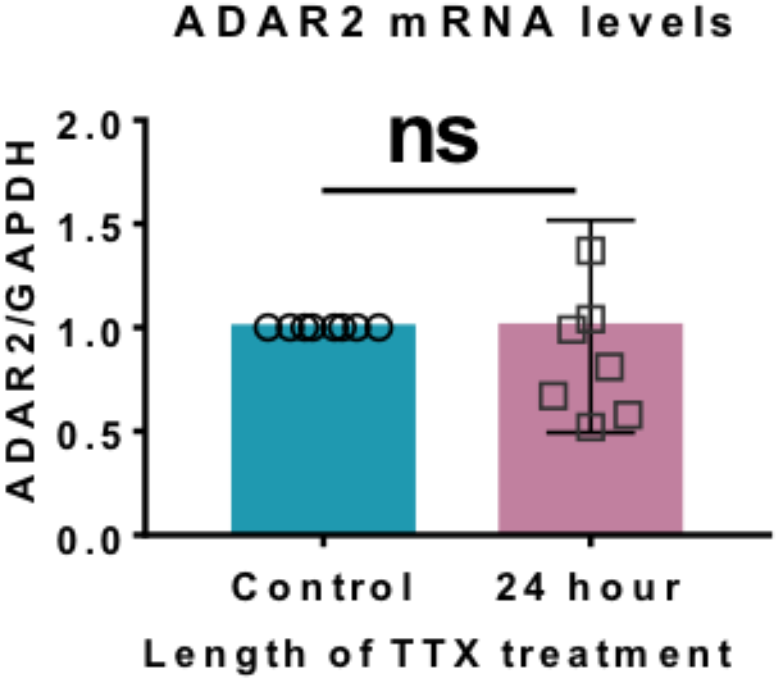
RT-qPCR analysis of mRNA levels of ADAR2 post TTX treatment showing no changes in the ADAR2 mature mRNA transcripts. Statistical analysis: Unpaired t-test; ns>0.05.

**Supplementary Figure 4:**
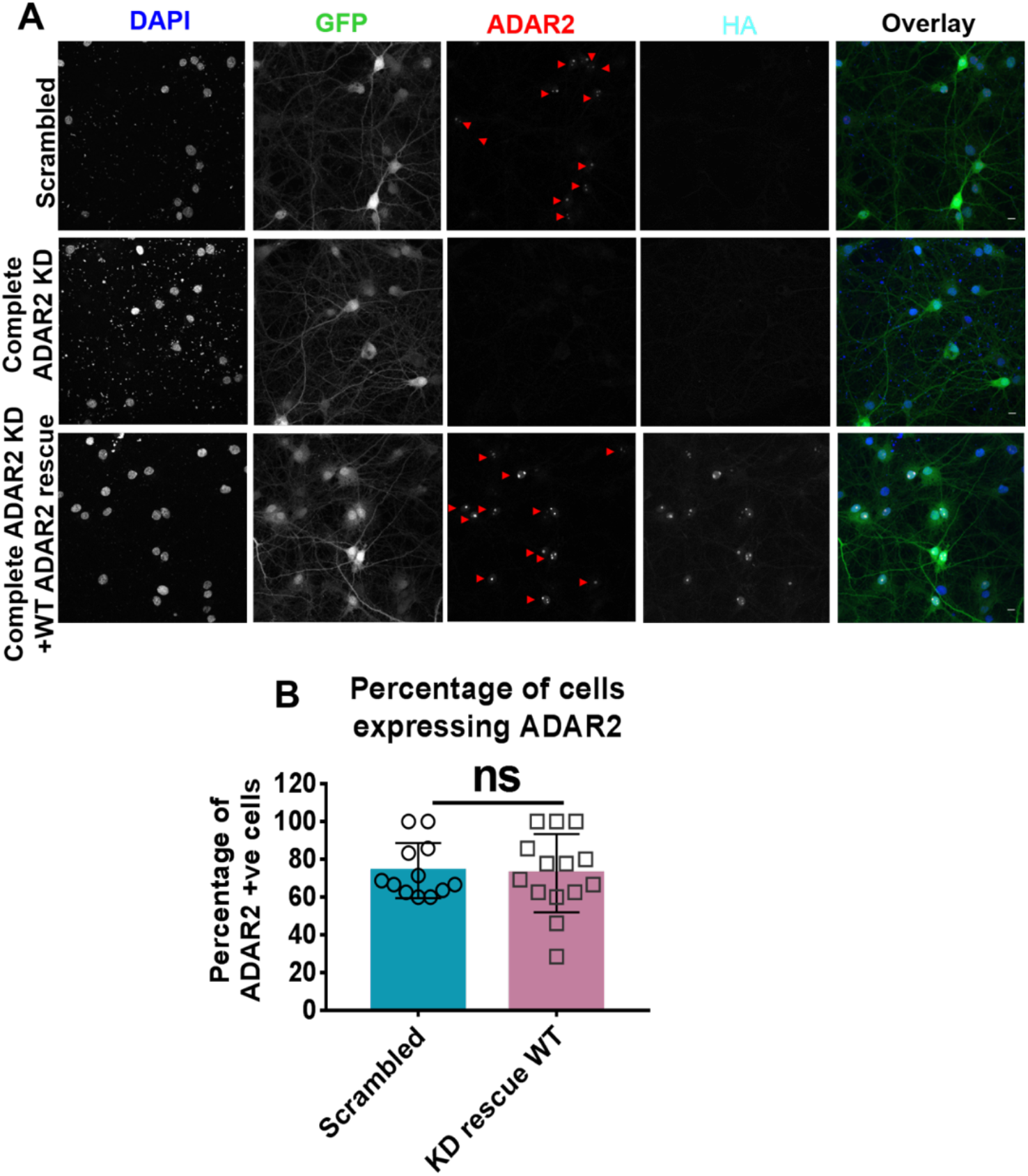
Confocal imaging showing successful knockdown and rescue of endogenous ADAR2 with HA tagged WT ADAR2. A. Representative confocal images showing nucleus imaged for DAPI (nucleus), GFP to represent the cells infected with ADAR2 knockdown, anti-ADAR2 to show successful rescue of ADAR2 in the same cells infected with the knockdown and anti-HA to show the rescues were HA tagged. B. Quantification showing the percentage of the cells expressing the rescue is comparative to the scrambled. N=4 independent dissections and n=12-14 fields of view. Statistical Analysis: Unpaired t-test; ns>0.05.

**Supplementary Figure 5:**
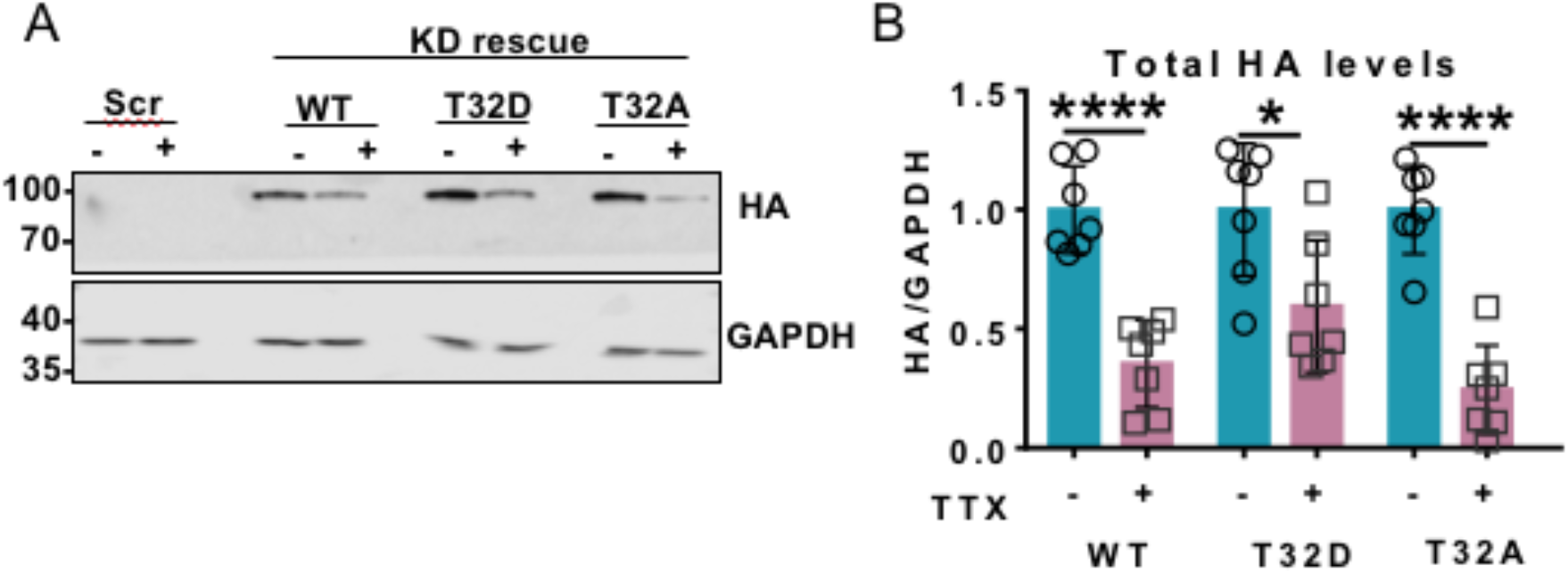
HA-tagged WT, phosphonull (T32A) or phosphomimetic (T32D) ADAR2 are all equally sensitive to the 24 h TTX treatment. A. Representative western blots of HA and GAPDH in cells infected with knockdown and rescued with HA tagged version of WT or T32A or T32D version of ADAR2. B. Quantification of (A) HA signals were normalised to GAPDH. Each TTX treated condition were normalised to their respective non-treated controls. Statistical Analysis: Unpaired t-test; *<0.05, ****<0.0001.

**Supplementary Figure 6:**
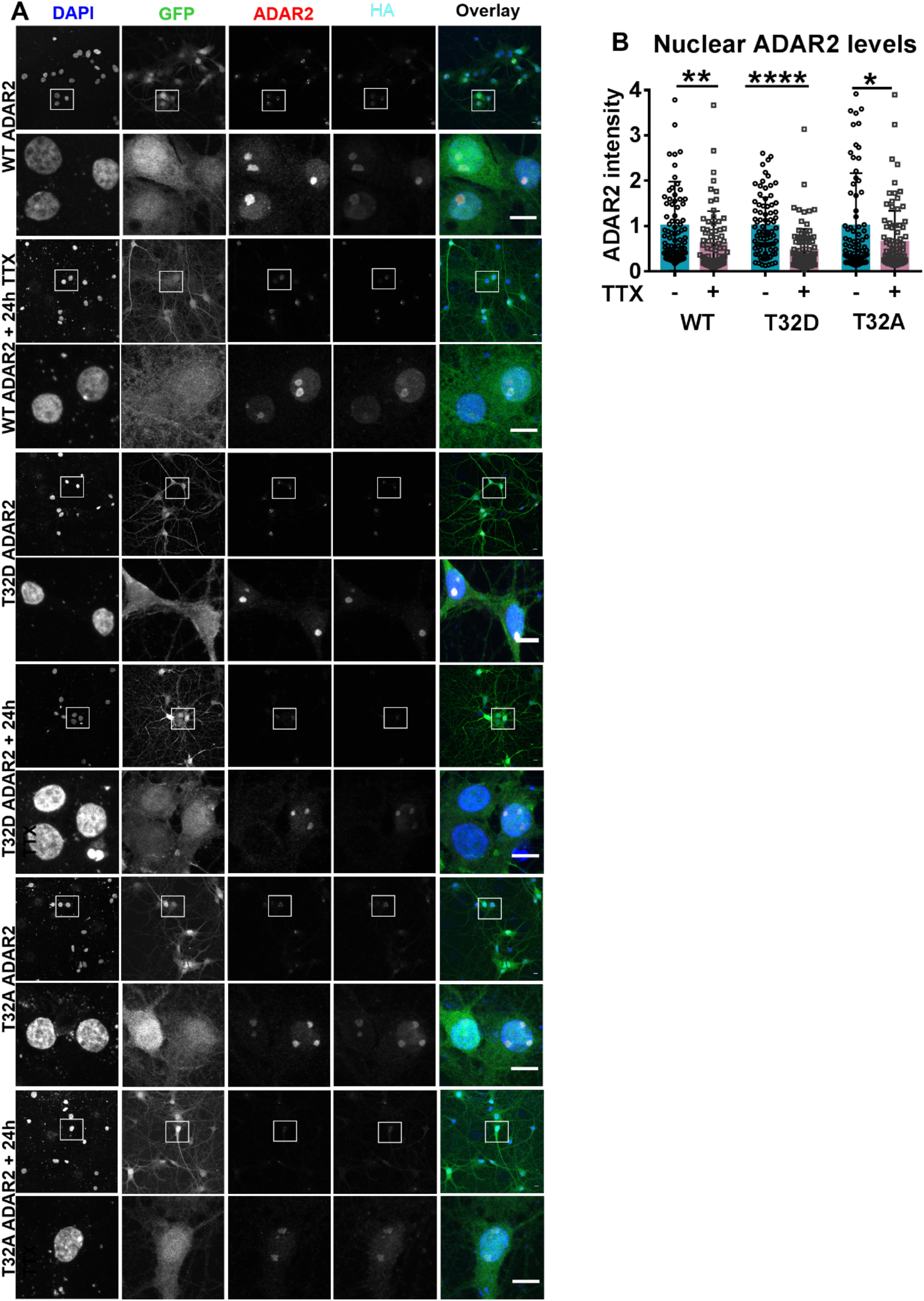
Confocal imaging showing the HA tagged WT, T32A and T32D ADAR2 are localised in the nucleus and sensitive to TTX. A. Representative images of hippocampal neurones imaged for DAPI (blue), GFP (green; infected cells), ADAR2 (red) and HA (tag) (cyan) following ADAR knockdown and rescue. The lower panels for each group show enlargements of the areas indicated by the box. B. Quantification of ADAR2 intensity in the nucleus of the images in A). TTX treated conditions were normalised to their respective non-treated controls. Statistical Analysis: Unpaired t-test; *<0.05, **<0.01, ****<0.0001. N=3 independent dissections and n=81-92 cells.

